# ProTECT – Prediction of T-cell Epitopes for Cancer Therapy

**DOI:** 10.1101/696526

**Authors:** Arjun A. Rao, Ada A. Madejska, Jacob Pfeil, Benedict Paten, Sofie R. Salama, David Haussler

**Affiliations:** Department of Biomolecular Engineering, University of California, Santa Cruz, Santa Cruz, CA, USA; Computational Genomics Lab, University of California, Santa Cruz, Santa Cruz, CA, USA; UC Santa Cruz Genomics Institute, University of California, Santa Cruz, CA 95064, USA; Department of Molecular, Cell, and Developmental Biology, University of California, Santa Cruz, Santa Cruz, CA, USA; Howard Hughes Medical Institute, University of California, Santa Cruz, CA 96064, USA

**Keywords:** cancer, neoepitope, neoantigen, prediction, automated, vaccine

## Abstract

Somatic mutations in cancers affecting protein coding genes can give rise to potentially therapeutic neoepitopes. These neoepitopes can guide Adoptive Cell Therapies (ACTs) and Peptide Vaccines (PVs) to selectively target tumor cells using autologous patient cytotoxic T-cells. Currently, researchers have to independently align their data, call somatic mutations and haplotype the patient’s HLA to use existing neoepitope prediction tools. We present ProTECT, a fully automated, reproducible, scalable, and efficient end-to-end analysis pipeline to identify and rank therapeutically relevant tumor neoepitopes in terms of immunogenicity starting directly from raw patient sequencing data, or from pre-processed data. The ProTECT pipeline encompasses alignment, HLA haplotyping, mutation calling (single nucleotide variants, short insertions and deletions, and gene fusions), peptide:MHC (pMHC) binding prediction, and ranking of final candidates. We demonstrate ProTECT on 326 samples from the TCGA Prostate Adenocarcinoma cohort, and compare it with published tools. ProTECT can be run on a standalone computer, a local cluster, or on a compute cloud using a Mesos backend. ProTECT is highly scalable and can process TCGA data in under 30 minutes per sample when run in large batches. ProTECT is freely available at https://www.github.com/BD2KGenomics/protect.

## 1 Introduction

Tumor recognition by the adaptive immune system has been described in the literature as early as the 1980s. In 1987, Muul et al. described tumor infiltrating lymphocytes in a cohort of 6 melanoma samples that showed high cytotoxicity towards fresh, autologous melanoma tumor cells (1). However, at the time, T-cell responses were observed to be short lived, often lasting only a few days. Later studies showed that tumors were capable of suppressing immune responses via different methods (2–5).

Checkpoint blockade therapy has seen a great increase in interest in the past few years with numerous drugs being approved by the FDA for clinical treatment (6–8). Prevention of PD-1:PD-L1 (9) and CTLA-4:B7.1/2 (10) binding via monoclonal antibodies re-enables the immune attack against the tumor, however it can leave the patient open to development of autoimmunity or other toxicities associated with unchecked immune action (11, 12). The mutational load of a tumor (or Tumor Mutational Burden) is a good predictor of response to checkpoint therapy (13, 14). The implication of aberrations in DNA Mismatch repair genes in impairment of tumor growth (15) suggest this effect is due to tumor “neoantigens” that act as markers for immune targeting.

Adoptive cell therapies use T-cells specifically targeted against the tumor to reduce the collateral damage associated with conventional therapies. Chimeric Antigen Receptor (CAR) cells use cell surface receptors with an antibody as the recognition domain and the downstream effector domains of a T-Cell receptor to elicit a T-cell response upon cognate epitope recognition (16). This method has been well studied in B-Cell lymphomas and leukemias and shows great promise due to the tissue-specific expression of targets like CD19, exclusively expressed on B-cells (17, 18). There are several attempts to apply CAR therapy to other cancers (19, 20).

Tumor Infiltrating Lymphocytes from patient tumors can be activated and expanded in-vitro using minced autologous tumor or experimentally primed autologous dendritic cells (21, 22). These cells selectively target cell-surface MHC-presented antigen produced by the tumor. Peptide vaccines attempt to produce the same result by stimulating dendritic cells in-vivo via synthetically produced peptides delivered subcutaneously to the patient. Experimentally primed Dendritic cells and Peptide vaccine therapies require prior knowledge of the mutations in the tumor in order to identify the potentially targetable sequence.

Bioinformatic analysis of tumor sequencing data can aid in the selection of neoepitopes to target in vaccine and adoptive immune system-based cancer therapies. PVAC-Seq (23) is an automated pipeline that identifies neoepitopes generated from a pre-computed, VEP-annotated (24) VCF file run with specialized plug-ins that incorporate wildtype and mutant protein sequence. INTEGRATE-Neo (25) identifies neoepitopes from fusion genes provided in a pre-computed BEDPE file. These tools all require a user to previously align the sequencing data to a reference of choice and call variants before following the same logical paradigm of identifying mutant peptides and predicting pMHC affinity binding (often via netMHC (26)). The pipelines differ in their degree of automation, input mutation type and annotation, and presence or absence of a ranking schema, however neither of them fully automates the pipeline from end-to-end, beginning at the raw fastq files emitted by the sequencer from DNA and RNA sequencing.

We describe ProTECT, a fully automated tool for the Prediction of T-cell Epitopes for Cancer Therapy. We previously demonstrated the utility of ProTECT, using an early version to analyze externally called Single Nucleotide Variants (SNVs) in a neuroblastoma cohort. There we identified a potentially therapeutic neoepitope from the ALK:R1275Q hotspot mutation (27). The full ProTECT codebase, reported here, is completely self-contained. It accepts an input trio of sequencing data from a patient consisting of the paired tumor and normal DNA, and the tumor RNA reads in the fastq format and processes the data from end-to-end including alignment, in-silico HLA haplotyping, expression profiling, mutation calling and neoepitope prediction.

We demonstrate the scalability and utility of ProTECT, and evaluate its performance, using publicly available data. We use the 326 samples from The Cancer Genome Atlas (TCGA) Prostate Adenocarcinoma (PRAD) cohort (28) with trios of genomic data (Tumor DNA, Normal Dna, and Tumor RNA), augmenting these data with 8 previously published clinical melanoma samples (29). The TCGA PRAD data set was previously evaluated for fusion-gene-derived epitopes using INTEGRATE-Neo (25) and the melanoma samples were previously analyzed by PVAC-Seq (23) We compared ProTECT’s performance to the performance of these other tools. The TCGA PRAD cohort has an average of 21.5 exonic mutations per sample (30) and 31% of all samples are predicted to contain a fusion transcript (31). The melanoma dataset was reported to have between 219 and 598 missense exonic mutations per sample. We show that ProTECT outperforms both PVAC-Seq and INTEGRATE-Neo by identifying the expected peptides with a low false positive rate, in addition to other events missed by the original methods.

## 2 Materials and methods

### 2.1 Procurement of Input Data

Genomic Trio (Tumor DNA, Normal DNA, and Tumor RNA) BAM files containing sequences from 326 samples in the Cancer Genome Atlas (TCGA) Prostate Adenocarcinoma (PRAD) cohort were downloaded from the Genomics Data Commons (GDC) at the National Cancer Institute using the GDC data transfer tool. The downloaded BAM files were converted back to FASTQ format, as would be produced by direct sequencing, using the SamToFastq module from Picard version 1.125^1^. MHCI haplotype calls using POLYSOLVER (32) for all samples for benchmarking were obtained and used with the permission of Dr. Catherine Wu.

Genomic Trios from 3 additional samples (Mel-21, Mel-38, Mel-218) were downloaded from the NCBI short read archive (SRA) (33) via Bioproject PRJNA278450/dbGaP accession phs001005. These patients were diagnosed with stage III resected cutaneous melanoma and had all previously received ipilimumab. Data from seven A*02:01 restricted vaccines tested for each patient were obtained from the supplementary information of the original manuscript (29).

The input data for the INTEGRATE-Neo comparison included haplotype and fusion calls from 240 samples in the supplementary data of the INTEGRATE-Neo paper. The fusions from supplementary Excel sheet 1 were parsed into individual BEDPE format files and the epitopes from sheet 3 were extracted into individual haplotype list files with one MHC allele per line.

Indexes for the various tools were generated using the GRCh38 (hg38) reference sequence obtained from the UCSC genome browser (34). GENCODE (35) v25 was chosen as the reference annotation and was used to filter the background SNV databases and SNPEff. Every generic hg38 index used in the analysis is available in our AWS S3 bucket ‘protect-data’ under the folders ‘hg38_references’. A detailed list of commands used to create the various indexes is available in the same bucket in the ‘README’.

### 2.2 Compute resources utilized

All TCGA-related analyses were conducted on a Mesos (36) cluster with one leader (12 cpus, 62GB RAM, 500GB Local disk) and eight identical agents (56 cpus, 250GB RAM, 1.8TB local disk).

The Melanoma data was analysed on the Amazon Web Services EC2 and the data was stored securely using SSE-C encryption on S3.

### 2.3 326-sample PRAD compute

The 326 samples were run in batches of 1, 2, 5, 10, 20, or 50 samples in order to gauge the efficiency and scalability of the pipeline engine, Toil (37). Each batch size was run 5 times with unique samples to normalize the runtime information. The configuration file for each run was generated from a template containing all the required tool options and paths to the input reference files on the NFS storage server. Each batch was run once on the mesos cluster using all nodes and an NFS-based Toil file job store to save the state of the pipeline. The five single-sample batches were also run separately without mesos on individual nodes of the cluster using an NFS-based Toil file job store to document the time taken per sample on a single machine.

### 2.4 Comparison with PVAC-Seq

To compare our results with PVAC-Seq, we ran ProTECT on the input samples on AWS EC2 using an S3-based cloud job store. The input configuration for the run included paths to hg38-mapped reference files from our S3 bucket ‘protect-data’ and paths to the input FASTQ files in another secure bucket. The results were stored on S3 in the same bucket as the input. This analysis was conducted consistent with the mandatory cloud data use limitations on the input dataset.

### 2.5 Comparison with INTEGRATE-Neo

To compare our results with INTEGRATE-Neo, we parsed the data from the manuscript supplement into files acceptable by ProTECT via a python script. The initial input configuration file consisted of links to the fusion BEDPE format file for each of 240 samples, along with haplotype and expression data called from the ProTECT 326 sample run. The final analysis included fusion and inferred haplotype calls for 83 samples from INTEGRATE-Neo along with ProTECT expression estimates. All ProTECT runs were conducted on the mesos cluster.

## 3 Pipeline specifics

ProTECT consists of 8 major sections: sequence alignment, haplotyping, expression profiling, mutation calling, mutation translation, MHC:peptide binding prediction, neoepitope ranking, and reporting. Figure 1 shows the schema for the run. Every tool used in the pipeline was hand-picked from industry-standard choices and literature reviews. Some aspects of the pipeline, notably Transgene and Rankboost, were developed in-house due to a lack of publicly available alternatives.

**Figure 1.**
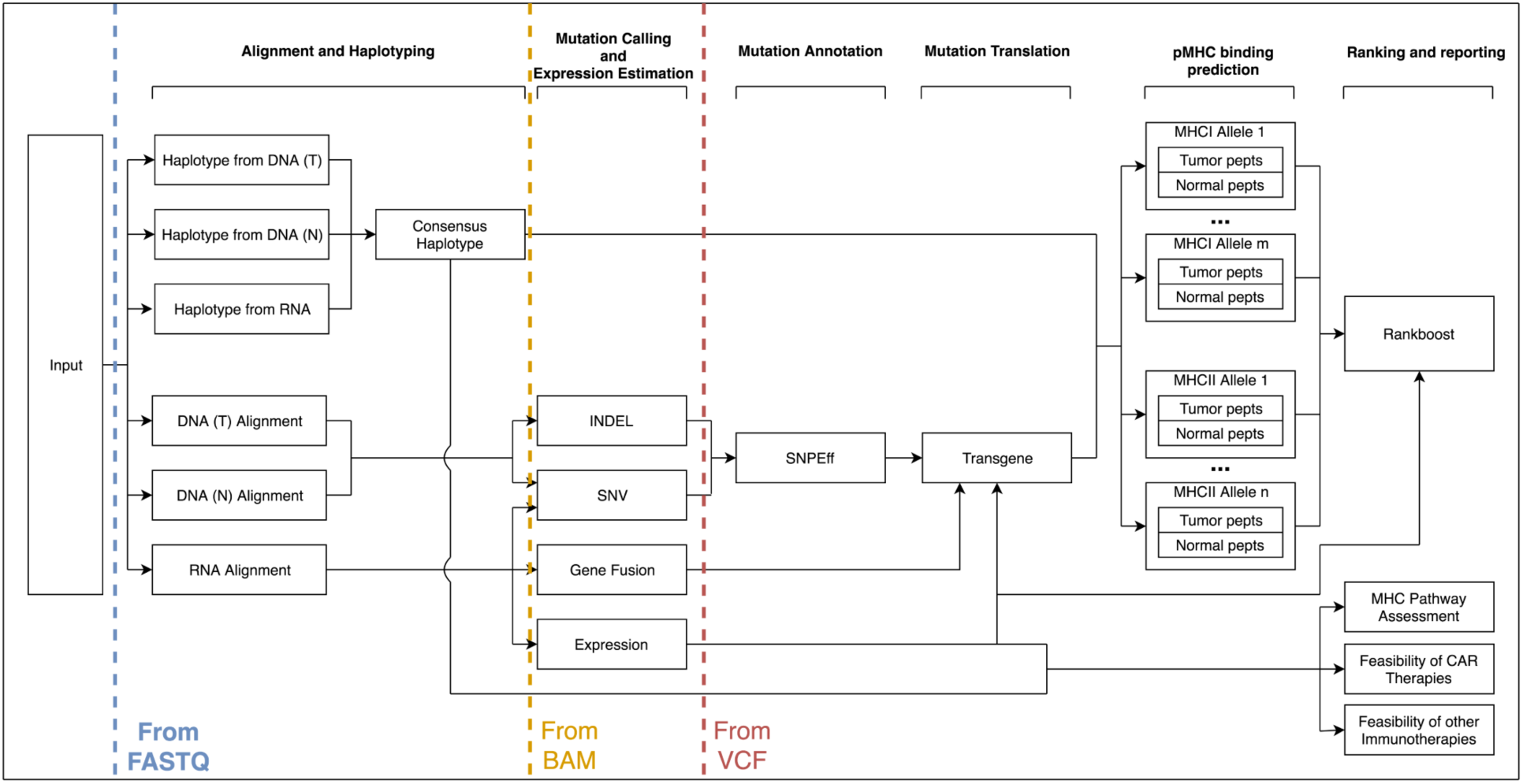
A schematic description of the ProTECT workflow. ProTECT can process FASTQs all the way through the prediction of ImmunoActive Regions, including alignment, HLA Haplotyping, variant calling, expression estimation, mutation translation, and pMHC binding affinity prediction. ProTECT also allows users to provide pre-computed inputs for various steps instead.

The entire analysis from end-to-end is built to process data against the same reference sequence and annotation. The user provides links to the properly generated indexes for each tool in the pipeline. We provide Gencode (35) version 19 annotated references for hg19 and Gencode version 25 annotated references for hg38 on our public AWS S3 bucket “protect-data”. The input for a protect run is a single configuration file that lists input files for each patient that will be processed, and all the options and links to indexes that will be used during the run.

## 3.1 Sequence Alignment

DNA sequence alignment is carried out using the Burrows-Wheeler aligner (BWA) (38). The reads are aligned with BWA-MEM to the provided BWA reference using default parameters. The SAM file produced upon alignment is processed to properly format the SAM header, and is then converted to a coordinate-sorted BAM file with a corresponding index.

RNA sequence alignment is carried out using the ultra-fast aligner, STAR (39). The parameters for the run are optimized for fusion detection via STAR-fusion (40).

Alternatively, ProTECT accepts pre-aligned BAM files as an input if the MHC haplotype is provided as well. ProTECT assumes that the user has aligned the DNA and RNA using the same reference genome with the same genomic annotation.

### 3.2 Haplotyping

The HLA Haplotype of the patient is predicted using PHLAT (41). The haplotype is predicted using each input source of information (normal and tumor DNA, tumor RNA) and the consensus haplotype is generated based on agreement between the three haplotype predictions. Due to limitations in the tool, we only proceed with HLA-A, HLA-B and HLA-C for MHCI, and HLA-DPA/B and HLA-DRB for MHCII.

### 3.3 Expression profiling

The gene-level and isoform-level expression is estimated using RSEM (42) with default parameters.

### 3.4 Mutation Calling

Single Nucleotide Variants (SNVs) are predicted on a per-chromosome basis using 5 separate mutation prediction algorithms: MuTECT (43), MuSE (44), RADIA (45), Somatic Sniper (46), and Strelka (47). The choice of mutation callers was guided by the results from the ICGC DREAM mutation calling challenge (48). All called mutations are merged into a common file and only events supported by 2 or more predictors advance to the translation step. Strelka additionally produces a callout for short insertions and deletions (INDELs). These are also used to identify neoepitopes.

Fusion calling occurs using STAR-Fusion (40) with default parameters. Candidate fusions are annotated using Fusion-Inspector^2^ along with an optional assembly step using Trinity (49).

### 3.5 Mutation Translation

SNVs and INDELs are annotated using SNPEff (50). Mutations identified in coding regions of the genome are processed using an in-house translation tool, Transgene^3^. Transgene filters the input SNPEff-produced VCF file to exclude non-expressed calls based on the gene expression data obtained in the previous step. SNVs and in-frame INDELs are directly injected into the amino acid chain to produce the mutant sequence. Frameshift INDELs are translated downstream of the mutation event till a stop codon is found (or a user-defined threshold is reached). Events lying within 27, 30, and 45 bp of each other (for 9-mer-, 10-mer- and 15-mer-containing peptides respectively) are chained together into an “Immunoactive region” (IAR), or a region that will potentially produce an immunogenic peptide. Separate mutation events that are combined into a single immunoactive region are phased using the RNA-Seq data to ensure that they truly are co-expressed on the same haplotype. Fusion IARs are generated using the breakpoints present in the input BEDPE file. TransGene uses provided junction sequences or infers them from the input annotation file. The predicted IAR contains (n-1)*3 bp on either side of the fusion junction from each donor for each n in 9-, 10-, and 15mer.

### 3.6 MHC:Peptide binding prediction

The predicted neoepitopes are assayed against each of the MHCI (9- and 10-mers) and MHCII (15-mers) predicted to be in the patient’s HLA haplotype using the IEDB MHCI and MHCII binding predictions tools.

The IEDB tools run a panel of methods (51–57) on each input query (input peptide fastq + MHC allele) and provides a consensus “percentile rank” that describes on average, how well each peptide is predicted to bind against a background set of 100,000 UniPROT derived peptides. Calls predicted to bind within the top 5% of all binders are selected for further study. The normal, unmutated (“wildtype”) counterpart peptide for each selected neoepitope is then also assayed against the MHC(s) identified to determine how well it binds, so that this can be compared to the binding affinity of the mutant version.

### 3.7 Neo-epitope ranking

Neoepitope:MHC calls are consolidated by the candidate IAR of origin. IARs are ranked using a boosting strategy (Supplementary Methods) that rewards candidates satisfying certain biologically relevant criteria including the number of calls originating from the IAR, the promiscuity of the region (i.e. the number of MHCs stimulated by peptides from the region), the combined expression of the isoforms displaying a neoepitope-generating mutation, the number of neoepitopes in the region predicted to bind to an MHC better than their wildtype counterpart, and the number of events where a 10-mer and 9-mer subsequence of it both bind well to an MHC (only done for MHCI). The user can specify the weights for each of these criteria if they do not want to use the defaults. The user can specify how much a candidate is allowed to be boosted up within a range of 0% to 55% (allowing the #1 position to be replaced by a better candidate). The algorithm conducts 3 iterations over the table of candidates to provide a final ranked list of epitopes in the sample. We ran our samples prioritizing overlap and promiscuity (0.68 and 0.32 respectively) for MHCI calls and set each covariate to 0.2 (equally important) for MHCII calls.

## 4 Results and Discussion

We ran 3 experiments to demonstrate our pipeline. The first experiment was run on 326 samples from the TCGA PRAD cohort and highlights the scalability, efficiency and utility of ProTECT. We also identify recurrent IARs in the cohort (containing mutations that occurred in more than one case) suggesting possible public neoepitopes for PRAD. The second experiment compares ProTECT to the published SNV- and INDEL-based neoepitope prediction pipeline, PVAC-Seq. The third experiment compares ProTECT to the published fusion-based neoepitope predictor, INTEGRATE-Neo. In all experiments, ProTECT was run using a consensus of 2 out of 5 mutation callers (as described above) and using all TransGene fusion filters to remove inter-mitochondrial, inter-immunoglobulin, 5’ lincRNA, and transcriptomic readthrough events. Results were tabulated using a mix of python scripts and manual curation on a local machine.

### 4.1 326 Sample run

To describe the scalability, utility and efficiency of ProTECT, we ran ProTECT on a total of 326 genomic trios from the TCGA PRAD cohort. We called a median of 79.5 SNVs and INDELs, and 7 fusion genes per sample, and accepted 20 and 3 respectively for the production of IARs. We identified a median of 11 IARs per sample. Of the 326 samples, only 3 samples were predicted to have no IARs. These samples were observed to have no expressed non-synonymous mutations or filter-passing fusions. The entire metrics table is presented in Supplementary Table 1 and the results are submitted in Supplementary File 1.

**Table 1.** Recurrent TMPRSS2-ERG breakpoints in the cohort. IARs from 21:41498119-21:38445621 and 21:41507950-21:38445621 are recurrent suggesting their viability universal peptide vaccine candidates. We do not expect to see an IAR from fusions with 5’UTR breakpoints. **\*:** TransGene cannot handle *de novo* splice acceptors. **\*\***: An Epitope will exist where the TMPRSS2 reads into the intron of ERG. **\*\*\*:** A frameshift is seen on the ERG side of the fusion.

| Breakpoint | Count | 5' breakpoint | 3' breakpoint | Neoepitope Expected? | IAR | Count |
| --- | --- | --- | --- | --- | --- | --- |
| 21:41508081-21:38445621 | 122 | 5' UTR | Exon 2 | No | NA |  |
| 21:41498119-21:38445621 | 48 | Exon 2 | Exon 2 | Yes | DNSKMALNS EALSVVSED | 37 |
| 21:41507950-21:38445621 | 35 | Exon 1 | Exon 2 | Yes*** | SGCEERGAA GSLISCE | 22 |
| 21:41508081-21:38474121 | 18 | 5' UTR | Intron 1 | No | NA |  |
| 21:41506445-21:38445621 | 18 | Intron 1 | Exon 2 | Yes* | NA |  |
| 21:41508081-21:38584945 | 11 | 5' UTR | 5'UTR | No | NA |  |
| 21:41498119-21:38474121 | 7 | Exon 2 | Intron 1 | Yes** | DNSKMALNS LNSIDDAQL | 7 |
| 21:41508081-21:38423561 | 7 | 5' UTR | Exon 3 | No | NA |  |
| 21:41498119-21:38423561 | 4 | Exon 2 | Exon 3 | Yes*** | DNSKMALNS ELS | 1 |
| 21:41494356-21:38445621 | 3 | Exon 3 | Exon 2 | Yes*** | SPSGTVCTS RSLISCE | 3 |

Figure 2 shows the results from running ProTECT with different batch sizes on our local cluster (See section Compute resources utilized). As the number of samples increases, we see an expected increase in overall time, but the average time per sample decreases drastically. We processed samples from end-to-end at a rate of 24.6 minutes per sample when running in a batch of 50 samples.

**Figure 2.**
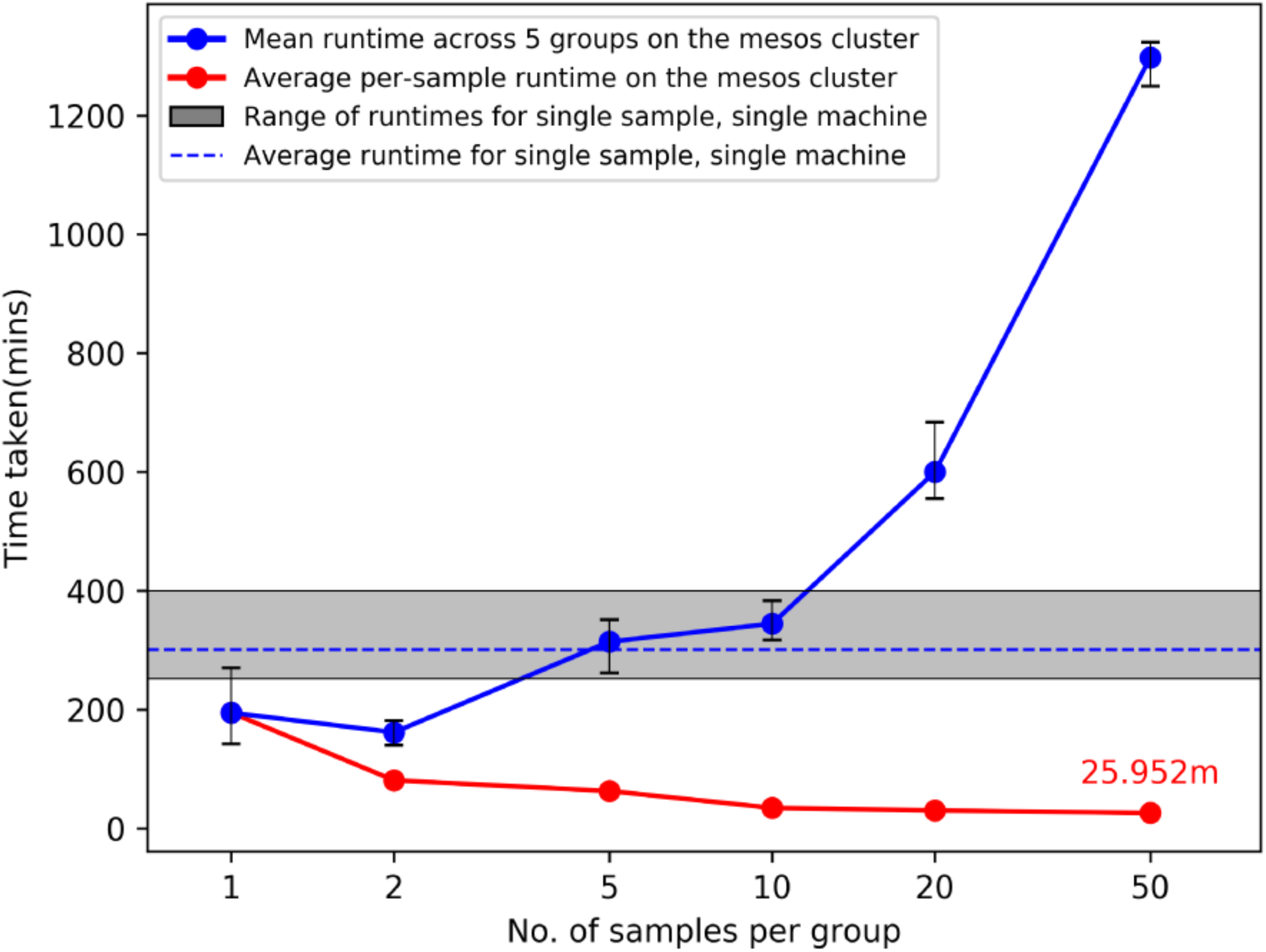
Average runtimes on our cluster when ProTECT is run in a batch of ‘n’ samples. Each batch of size ‘n’ is run with 5 unique sample sets and the range of runtimes is described by the whiskers at each datapoint. The grey bar describes the result of running ProTECT on a single sample on one machine. ProTECT takes considerably less time on average when run in a large group.

We detected the well-documented TMPRSS2-ERG fusion gene in 131 samples. We predicted at least one IAR each arising from 5 of the 10 unique breakpoints called (Table 1). Of the 5 breakpoints that do not result in an IAR, 4 of these breakpoints are located in the 5’ UTR of TMPRSS2 and will not result in a neoepitope. The 5th breakpoint has a 5’ intronic breakpoint and a 3’ exonic one, and the resulting neoepitope should contain the translated product from the last few bases of TMPRRS2 Exon 1 and the first bases after the *de novo* splice acceptor is reached in ERG. This case is not handled by TransGene at this time, and so no neoepitope call was made. One IAR of particular interest is DNSKMALNSEALSVVSED from the junction chr21:41498119-chr21:38445621, which is found in 37 of the 48 unique samples harboring that junction (11% of the entire cohort). Peptides from this IAR are predicted to bind well to HLA-A*02:01 (Allele Freq: 0.26) and HLA-C*07:01 (allele Freq: 0.17), alleles frequently seen in caucasian populations, which are highly represented in the TCGA cohort. Similarly, we predict SGCEERGAAGSLISCE from 22/35 samples with chr21:41507950-chr21:38445621, binding to C*07:01, C*04:01, B*44:02 (allele frequencies 0.14, 0.12, 0.08 respectively). The distributions of MHC alleles detected in patients harboring these events are shown in Supplementary Figures 1 and 2, respectively. These events are potentially viable candidates for public epitopes for patients with TMPRSS2-ERG, and could be pursued as vaccines for these cancers.

We detected a number of recurrent mutations in the SPOP gene concordant with previous reports (28, 58, 59). We detected 7 unique recurrent variants across 19 samples that map to 3 different amino acid positions in the SPOP protein, p.F133C/V/I/L, p.F102C/V, and p.W131G (Table 2). The mutation at position 133 might be of immunological interest since Leucine, Isoleucine and Valine have small hydrophobic side-chains and may stimulate the same TCR depending on pMHC binding. Samples with SPOP mutations lack ETV family fusions, suggesting that vaccine therapies against SPOP and the TMPRSS2-ERG fusion would target different populations of PRAD patients.

**Table 2.** Recurrent mutants in the SPOP gene target 3 codons. The F133V/C/I/L mutant may be of interest as a universal neoepitope due to the similar chemical properties of Leucine, Isoleucine and Valine.

| Variant | Count | Gene | Mutant | IAR | Frequency |
| --- | --- | --- | --- | --- | --- |
| chr17:49619064A>C | 5 | SPOP | p.F133V | RFVQGKDWG **V** KKFIRRDFL | 4 |
| chr17:49619063A>C | 3 | SPOP | p.F133C | RFVQGKDWG **C** KKFIRRDFL | 2 |
| chr17:49619064A>T | 2 | SPOP | p.F133I | RFVQGKDWG **I** KKFIRRDFL | 1 |
| chr17:49619062G>T | 2 | SPOP | p.F133L | RFVQGKDWG **L** KKFIRRDFL | 2 |
| chr17:49619281A>C | 2 | SPOP | p.F102C | CPKSEVRAK **C** KFSILNAKG | 2 |
| chr17:49619282A>C | 3 | SPOP | p.F102V | CPKSEVRAK **V** KFSILNAKG | 3 |
| chr17:49619070A>C | 2 | SPOP | p.W131G | AYRFVQGKD **G** GFKKFIRRD | 2 |

An important topic to highlight are HLA haplotypes called by PHLAT (41). We compared our results to the POLYSOLVER (32) calls and consistent with prior work (60), we see that PHLAT miscalled HLA-A*02:01 as HLA-A*01:81 in 33 samples. However, 29 of these samples are predicted to be homozygous HLA-A*02:01 by POLYSOLVER so the effect of this miscall will be to add information to the final ranked IARS from one additional allele. Since most IARs contain peptides predicted to bind to more than one allele, the noise produced by this artifact should not adversely affect the scores generated via the signal from calls against the correct partners. The remaining 4 samples were predicted to be heterozygous HLA-A*02:01/HLA-A*01:01 via POLYSOLVER and ProTECT identified these samples as HLA-*02:01/HLA-A*01:81. This is slightly worse than the first case since we’re completely lacking HLA-A*01:01 peptide binding affinity predictions for all these samples. Overall, 67.5% of all samples had perfectly concordant haplotypes with POLYSOLVER, 28.8% differed by 1 allele and 3.7% differed by 2 (Figure 3). A large chunk of the second group consists of the miscalls mentioned above. ProTECT allows users to provide pre-computed MHC haplotype calls if they trust another external caller more than PHLAT, or if they have haplotype information from another source.

**Figure 3.**
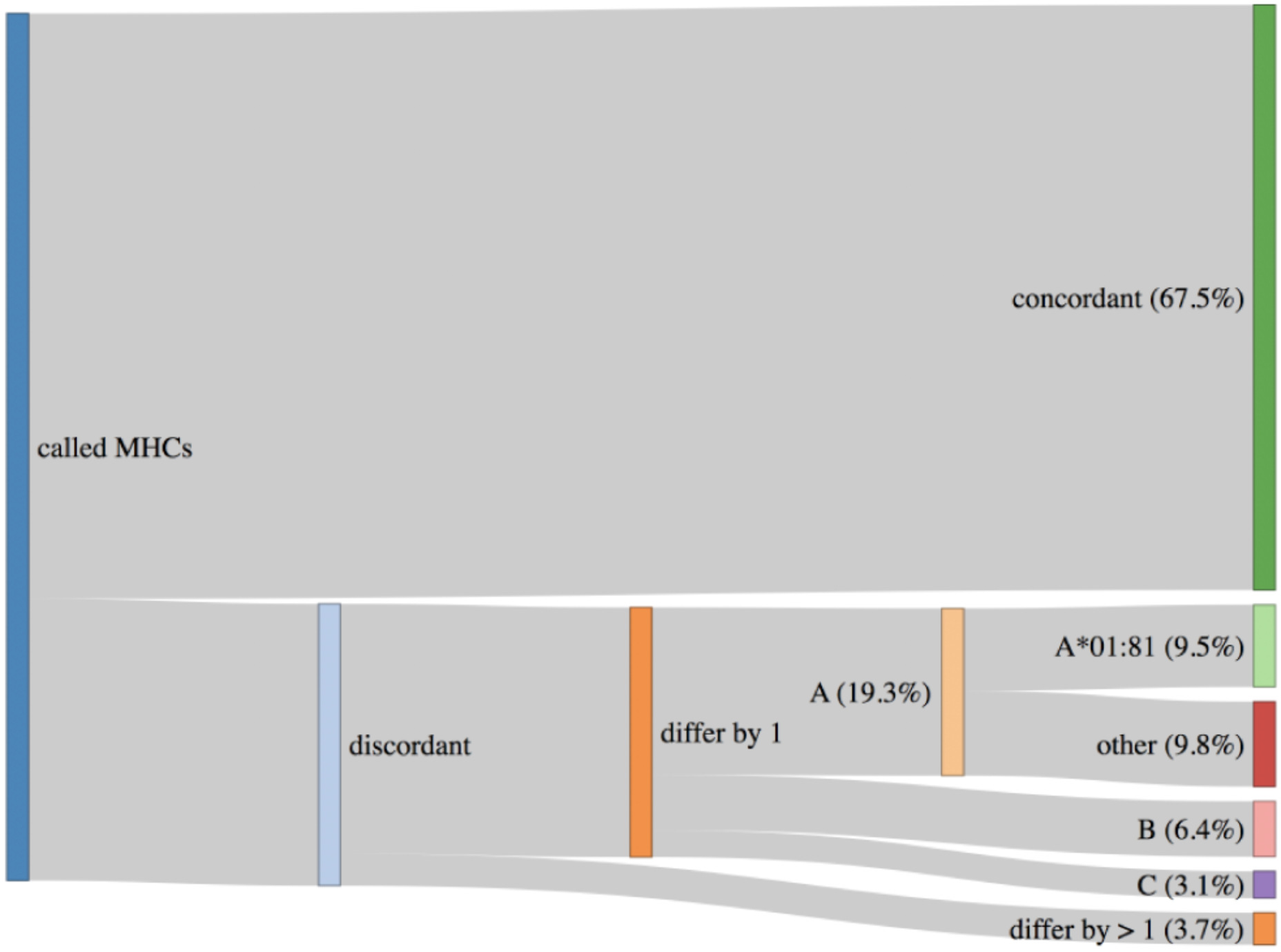
HLA Haplotypes called by ProTECT (using PHLAT) are fully concordant with POLYSOLVER haplotypes in only 67.5% of samples. 28.8% differ by 1 call and 3.7% by 2 calls. A majority of the miscalled HLA-A alleles are a documented PHLAT artifact.

### 4.2 Comparison with published callers

We ran ProTECT on the 8 melanoma samples from 3 patients (1 primary lymph node tumor each, and multiple metachronous tumors in 2 samples) (29) that were used to benchmark PVAC-Seq (23). Carreno et al. predicted 11-28 expressed, HLA-A*02:01 binding candidate peptides per sample and synthesized 7 unique peptide vaccines per patient based on presence of the mutants in the metachronous tumors, and assessed binding of the predicted peptide to HLA-A*02:01 in T2 assays. 3 peptides per patient were found to induce an immune reaction. ProTECT correctly identified the expected immunogenic mutations in every reported mutation:sample pair. In some cases, ProTECT even predicted the expected variant in a metachronous tumor where the original paper missed it (E.g. CDKN2A:E153K in the Lymph Node of Mel-21) (Table 3). Overall, ProTECT ranked IARS containing the validated variants relatively highly (in the top 15-20%) except in Mel218. We cannot definitively comment on the ranking in Mel218 since ProTECT considers every mutant and MHC allele in the MHC haplotype, while Carreno et al. only considered a curated list of peptides against HLA-A*02:01. In addition to the validated variants, we also provided a larger ranked set of possible candidates that broaden the spectrum of testable epitopes. The data for all 7 tested peptides is provided in Supplementary Table 2 and all neoepitopes predicted by ProTECT in Supplementary Table 3 and Supplementary File 2.

**Table 3.** ProTECT ranks on 8 Metachronous Tumors across 3 Melanoma patients [Hundal et.al]. Highlighted ranks describe instances where PVAC-Seq and ProTECT both call epitope. Green: Dominant epitope(existing immunity, neoantigen processed from endogenous protein), Orange: Subdominant epitope (immunity after vaccination, neoantigen processed from endogenous protein), Red: Cryptic epitope(immunity after vaccination, neoantigen not processed from endogenous protein).

| Sample and Source | Mel 21 |  |  |  | Mel 38 |  |  | Mel 218 |
| --- | --- | --- | --- | --- | --- | --- | --- | --- |
|  | LN | Skin (2012) | Skin (2013) |  | Abdominal Wall | Axilla LN | Breast | LN |
|  |  |  | RNA 1 | RNA 2 |  |  |  |  |
| Collection date | 1/30/2011 | 5/10/2012 | 6/6/2013 | 6/6/2013 | 4/16/2013 | 4/19/2012 | 2/14/2013 | 4/4/2005 |
| Total Variants | 1532 | 2140 | 1681 | 1679 | 1213 | 1121 | 1259 | 2176 |
| Actionable Variants | 332 | 400 | 393 | 391 | 219 | 216 | 224 | 449 |
| Total IARs | 105 | 137 | 114 | 116 | 73 | 80 | 86 | 155 |
| Vaccine Candidates with ProTECT ranks | NDC1:F169L (Reported as TMEM48 F169L) |  |  |  | SEC24A:P469L |  |  | EXOC8: Q656P |
|  | 4 | 5 | 10 | 3 | 8 | 9 | 2 | 152 |
|  | TKT:R438W |  |  |  | AKAP13:Q285K |  |  | PABPC1 :R520Q |
|  | 6 | 6 | 4 | 4 | 14 | 80 | 17 | 140 |
|  | CDKN2A:E153K |  |  |  | OR8B3:T190I |  |  | MRPS5: P59L |
|  | 21 | - | 18 | 17 | 11 | 13 | 14 | 26 |

We compared our fusion prediction accuracy with INTEGRATE-Neo (25). INTEGRATE-Neo was demonstrated on 321 samples from the TCGA PRAD cohort and at least 1 neoepitope was predicted from 161 samples. 240 of the 321 samples overlap with our 326 sample dataset and this subset was used for this experiment. None of the predicted neoepitopes in this study have been validated using any biological experiments. We first attempted to compare our fusions (called using STAR-Fusion) with the fusion calls generated from INTEGRATE (61). The overlap between the ProTECT and INTEGRATE calls was relatively low (595/1519, with 120 unique calls in ProTECT) but a large chunk of the non-overlapping calls were from events with 1 spanning read support in INTEGRATE(Supplementary Figure 3). We see a better overlap when we increase the minimum support to 2 (an internal metric within ProTECT), and also find that 44 events rejected for having 1 read support in INTEGRATE were detected by ProTECT with >1 read support. Some of the INTEGRATE-specific calls were picked up by ProTECT but filtered out as low quality events. We further noticed that the concordance between MHC haplotypes called by HLAMiner (62) (used by INTEGRATE-Neo), PHLAT (41) (Used by ProTECT), and POLYSOLVER (32) was very low (Supplementary Figure 4). 61 of the unique HLAMiner predictions across the cohort did not match any of the other two callers and 41 matched both. (Homozygous calls in a patient were treated as one call.) 2 alleles were shared exclusively between ProTECT and INTEGRATE and only 1 between INTEGRATE and POLYSOLVER. In order to conduct a more comparable analysis, we reran ProTECT with the INTEGRATE fusion calls and the MHC haplotypes from the INTEGRATE-Neo manuscript (182 neoepitopes from 720 fusions over 83 samples, Supplementary Figure 5). ProTECT rejected 100 of the 720 provided fusion events as transcriptional readthroughs (92) or for having a 5’ non-coding RNA partner (8). ProTECT correctly identified 139 neoepitopes as IARs and rejected the remaining for being in a rejected fusion (23), scoring below the 5% predicted binding score threshold (16), having a 5’ breakpoint in the UTR (3), or for having a 5’ non-coding partner (1) (Supplementary Table 4). On further inspection, we noticed that the 3 neoepitopes arising from the 5’ UTR breakpoints (TCGA-HC-7080, PRH1>>RP11-259O18.4 and PRH1>>M6PR) could have been detected if the 5’ partner had been annotated with a different gene (PRR4) at the same locus (Supplementary Figure 6), an issue arising due to the differing gene annotation gtfs used between the methods (Gencode v25 for ProTECT and Ensembl v85 for INTEGRATE). Interestingly, this type of event occurred in one other sample (TCGA-EJ-8474, C1QTNF3-AMACR>>NDUFAF2) however INTEGRATE called the overlapping call as well (AMACR>>NDUFAF2) and since the epitopes were identical from both, ProTECT picked them up under the correct call (Supplementary Figure 7). The full set of results from running ProTECT on 83 INTEGRATE-Neo inputs is provided in Supplementary File 3. Easing ProTECT’s 5% filter would increase the number of false positives called by too large a margin, so we stand by our decision to reject the 16 neoepitopes missed due to this filter. This experiment also highlights the modularity of protect, and it’s flexibility in accepting pre-computed inputs to run only the necessary steps to produce a ranked list of IARs.

### 4.3 Reproducibility

Every tool used the pipeline, from established aligners to the in-house script used to translate mutations, is wrapped in a Docker image (63). Docker allows a developer to wrap a piece of code, and any requirements, into an image that can be instantiated into a container on any other machine. The container is guaranteed to run in the same manner on any machine, under the same environmental constraints. This way, results from ProTECT runs on different machines will always be the same.

### 4.4 Automation, Scalability and Efficiency

ProTECT is built to be run end-to-end without any user intervention. ProTECT is written in the Toil framework and will attempt to run the pipeline on the given input samples in a resource-efficient manner. The pluggable backend Toil APIs allow protect ProTECT to run on a single machine, a grid engine cluster, or a mesos cluster setup on a local network or on AWS. Toil allows users to deploy scripts on Azure and the Google cloud as well, however ProTECT does not yet support these environments.

Users provide ProTECT a config file that details the input files, and the various indexes and versions of tools to use during the run. ProTECT downloads the files to a “file store” and then spawns a graph of jobs for each input sample culminating in a ranked list of epitopes. The nodes in the graph are tuned to request an appropriate number of CPUs (for multithreaded jobs), memory and disk space. Toil ensures that these jobs are parallelized to the maximum extent.

## 5 Conclusion

We have described an efficient, automated, and portable workflow for the prediction of Neoepitope candidates that can guide Vaccine-based or adoptive T-cell therapies. We have shown that ProTECT scales well on a parallel processing environment and shows great efficiency gains as the number of samples processed in a batch increases. On average, we processed a sample from end-to-end in 26.4 minutes when we ran 50 samples in a single batch on an 8-node cluster. We have shown that ProTECT is comparable to existing callers and improves on them by providing a ranked list of neoepitopes arising from SNVs, INDELs and fusion genes. None of the currently published pipelines give results for all three types of mutations. Positive results from a clinical trial were ranked highly in our results and we retrospectively picked up events missed by the caller (PVAC-Seq) used to guide the trial. We identified recurrent epitopes arising from the well-documented TMPRSS2-ERG fusion and these results suggest a peptide vaccine could be developed for one of the common breakpoints. While designed for use in the rapidly growing fields of cancer vaccines and Autologous T-cell therapies, ProTECT can also be used to understand the link between tumor mutational burden and response to checkpoint blockade therapies. It is our fervent hope that improvements in these fields will quickly establish neoepitope-targeted immunotherapies as standard-of-care for cancer treatment.

## 7 Author Contributions

AAR, SRS and DH contributed to the conception and design of the study; AAR developed the entire codebase available on github with significant contributions from JP; AAM was involved with analyses on the 326 sample run; AAR wrote the first draft of the manuscript; SRS, DH and AAR contributed to manuscript revision. DH, SRS and BP acquired funding for this work. All authors read and approved the submitted version.

## 9 Acknowledgements

The authors would like to acknowledge the contributions of Hannes Schmidt, Christopher Ketchum, Aisling O’Farrell, Swapnil Patil, Akul Goyal and Walter Shands towards the ProTECT source code, and Bala Ganeshan, Giridhar Basava, Hemant Trivedi of Stellus Technologies for the compute support that enabled the 326 TCGA sample run.

## Footnotes

1 Obtained from http://broadinstitute.github.io/picard/

2 Obtained from https://github.com/FusionInspector/

3 Hosted at https://github.com/arkal/Transgene

