## Supplementary Data and Figures for "ProTECT – Prediction of T-cell Epitopes for Cancer Therapy"

### Supplementary Material

#### 1 Supplementary Data

##### 1.1 Pseudocode for Peptide rank boosting

```
For i in 1, 2, 3
    ∀ candidate in candidates
      boost =  W_npa * boost_npa + \
               W_nph * boost_nph + \
               W_nMHC * boost_nMHC + \
               W_TPM * boost_TPM + \
               W_ovlp * boost_ovlp
      new_rank = old_rank * (1-boost)
```

Where W_x describes the weight for covariate x, boost_x describes the score for the candidate x from 0-1, npa=number of peptides constituting an IAR, nph=number of strongly binding peptides constituting the IAR, nMHC=number of MHCs predicted to recognize a neoepitope from this IAR, TPM=expression of the gene harboring the IAR, and ovlp=number of events where a 9-mer and 10-mer overlap and are predicted to bind to the same MHC (only valid for MHCI). The individual scores for each of the covariates was calculated empirically based on internal test data. The Ranking algorithm is available at https://github.com/arkal/rankboost.

### 2 Supplementary Figures, Tables, and Files

#### 2.1 Supplementary Figures

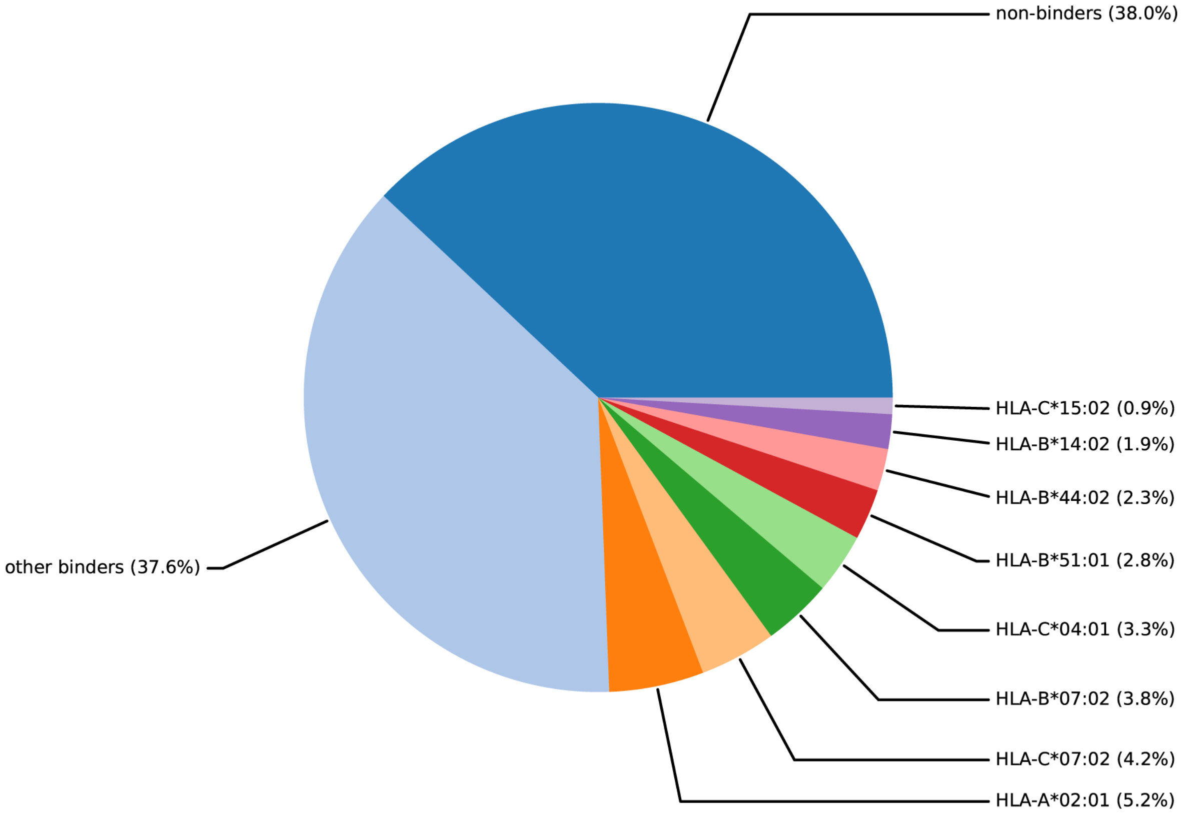


**Supplementary Figure 1.** Distribution of interesting MHC alleles called in samples harboring the TMPRSS2-ERG fusion IAR DNSKMALNSEALSVVSED at chr21:41498119-chr21:38445621

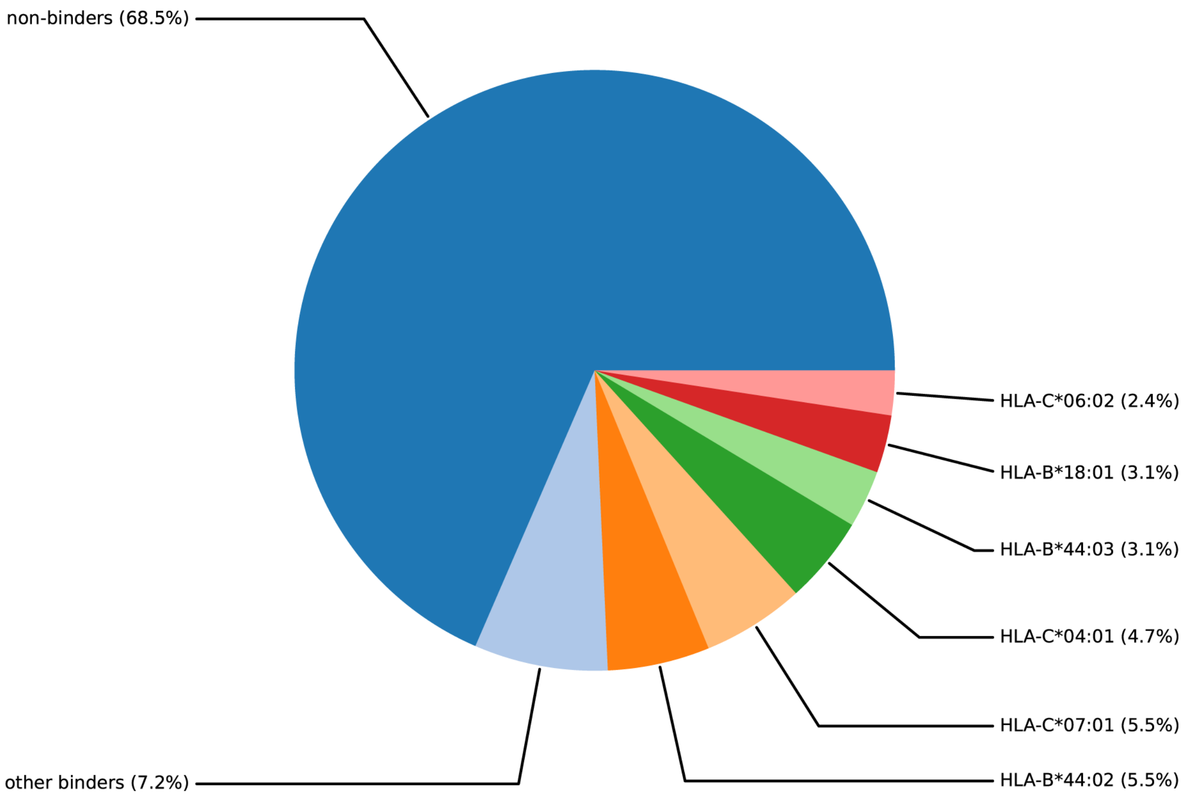


**Supplementary Figure 2.** Distribution of interesting MHC alleles called in samples harboring the TMPRSS2-ERG fusion IAR SGCEERGAAGSLISCE at chr21:41507950-chr21:38445621.

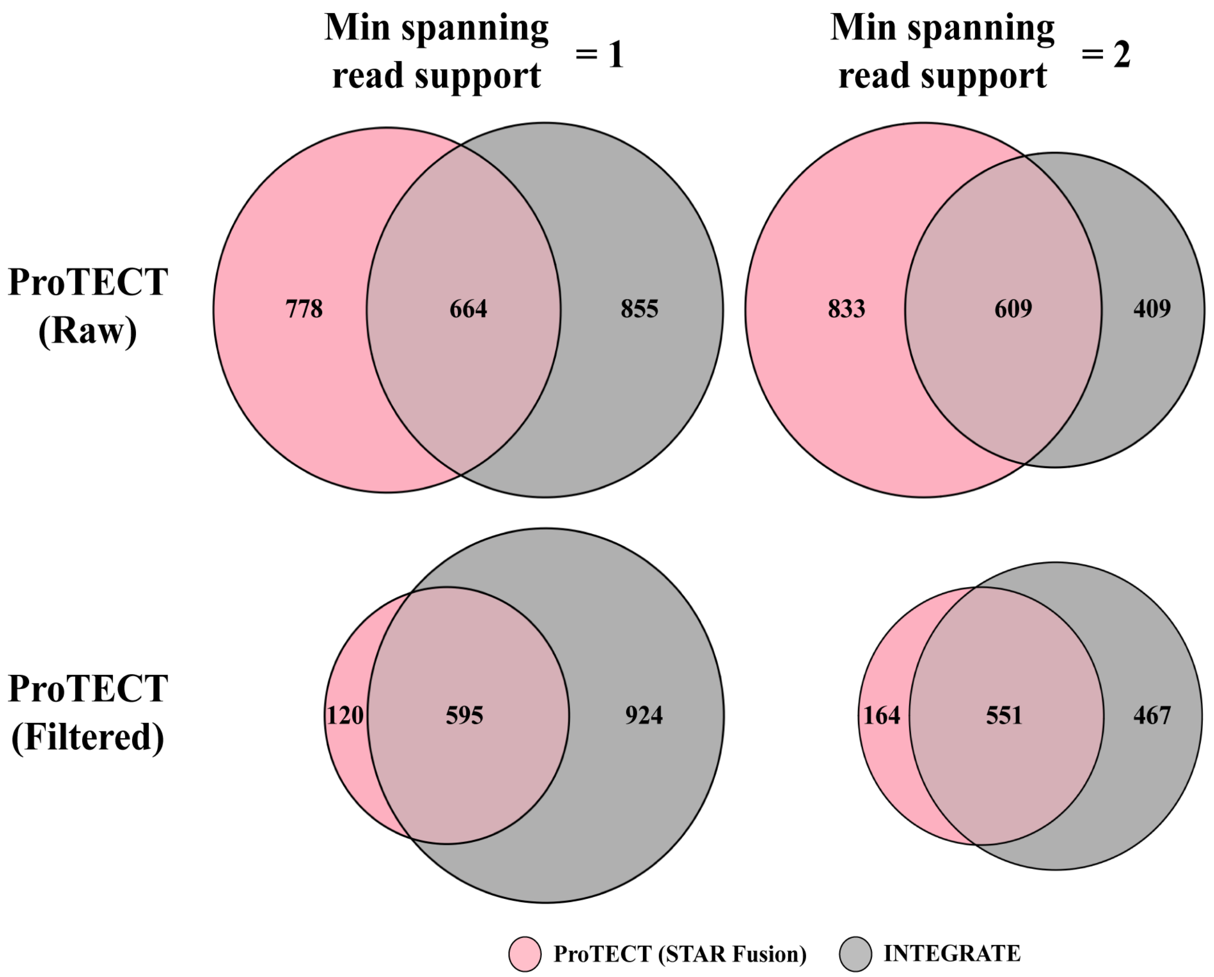


**Supplementary Figure 3.** Overlap between the fusion events called by ProTECT and INTEGRATE on a cohort of 240 prostate cancer samples. **Top**: There is low overlap between the raw ProTECT calls and INTEGRATE. Filtering INTEGRATE calls to retain only events with >= 2 supporting spanning reads (SSR) discards 33% of all calls but some of these events are called by ProTECT with >=2 SSR. **Bottom**: The filtered ProTECT results show better overlap with INTEGRATE calls using >=1 SSR and >=2 SSR. The number of ProTECT calls remains constant across rows since ProTECT uses a minimum of 2 SSR by default. The number of INTEGRATE calls remains constant across columns since there is no filtering applied to the INTEGRATE calls.

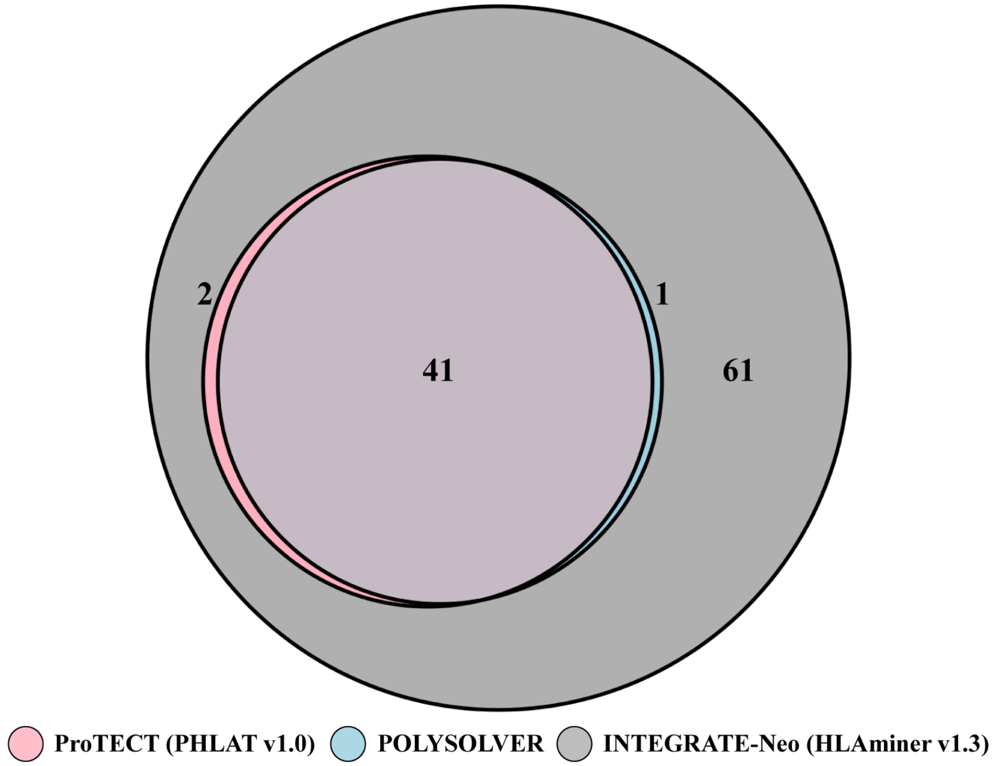


**Supplementary Figure 4.** HLA haplotypes called by HLAMiner in the INTEGRATE-Neo paper have very low overlap with ProTECT and POLYSOLVER.

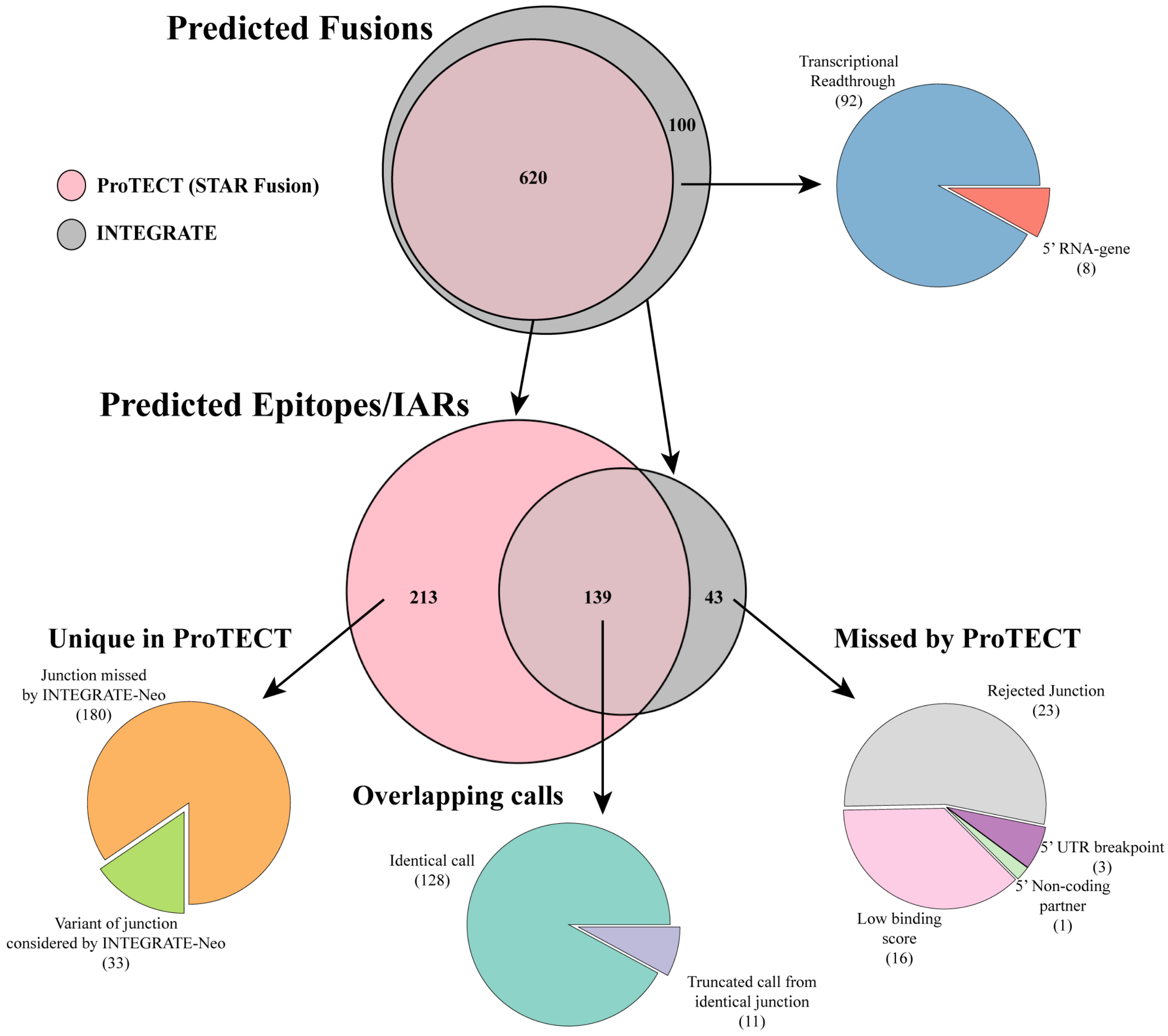


**Supplementary Figure 5:** A schematic comparing ProTECT and INTEGRATE-Neo, using the fusion and MHC calls from the INTEGRATE-Neo paper (for maximal comparability). Protect rejects 100/720 calls from 83 patients and calls 352 IARs from the rest. A majority (23/43) of the epitopes missed by ProTECT arose from the rejected junctions. Of the 139 overlapping IARs, a small fraction (11) were not identical to the INTEGRATE call, but had significant overlap with the INTEGRATE-Neo expected epitope.

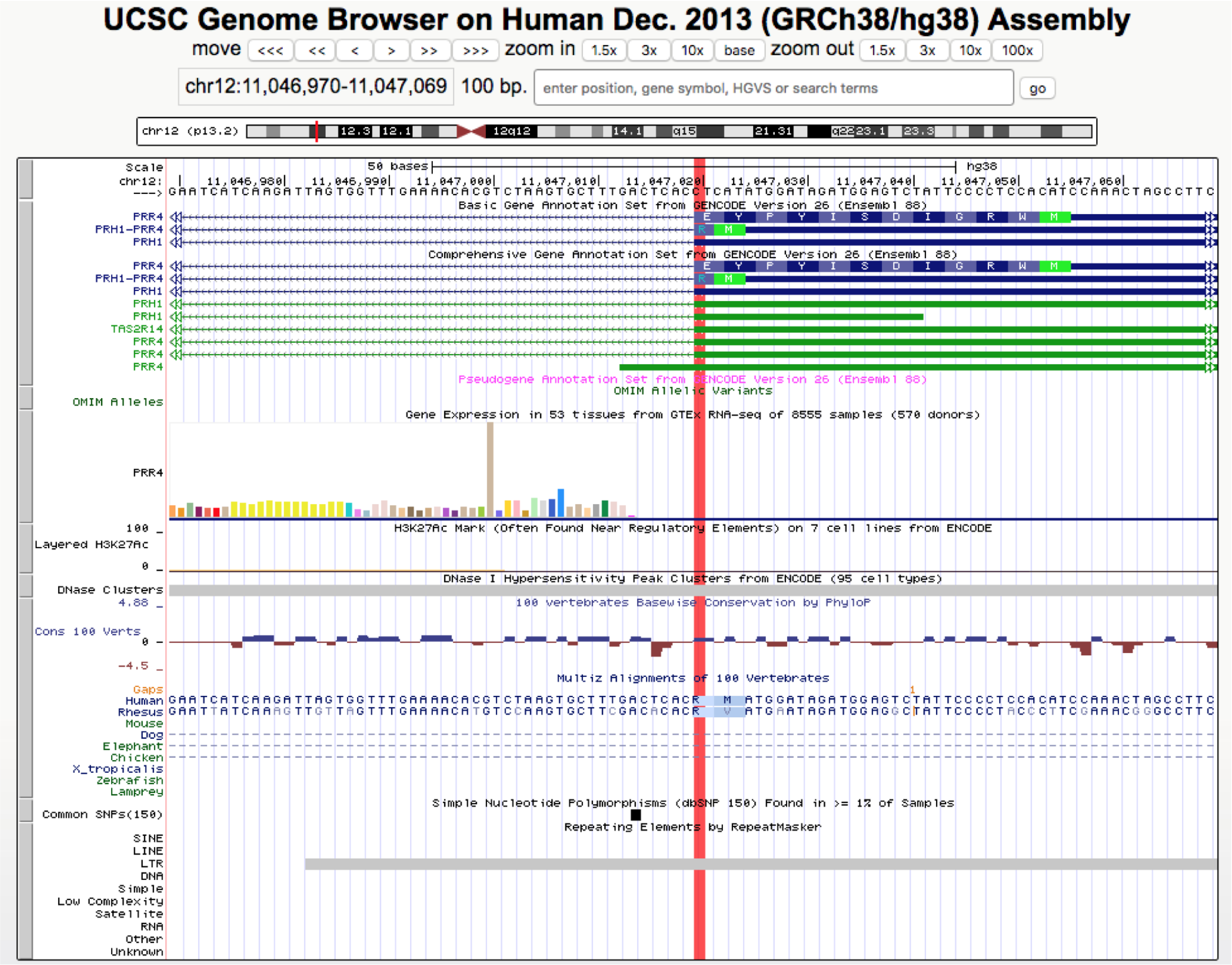


**Supplementary Figure 6.** A UCSC genome browser showing the 5' breakpoint (highlighted) for the two missed epitopes from the ENSG00000231887-ENSG00000003056 fusion. The 5' partner was reported as PRH1 but breakpoint is in the 5' UTR for PRH1. The overlapping PRR4 gene contains the breakpoint sequence predicted by INTEGRATE.

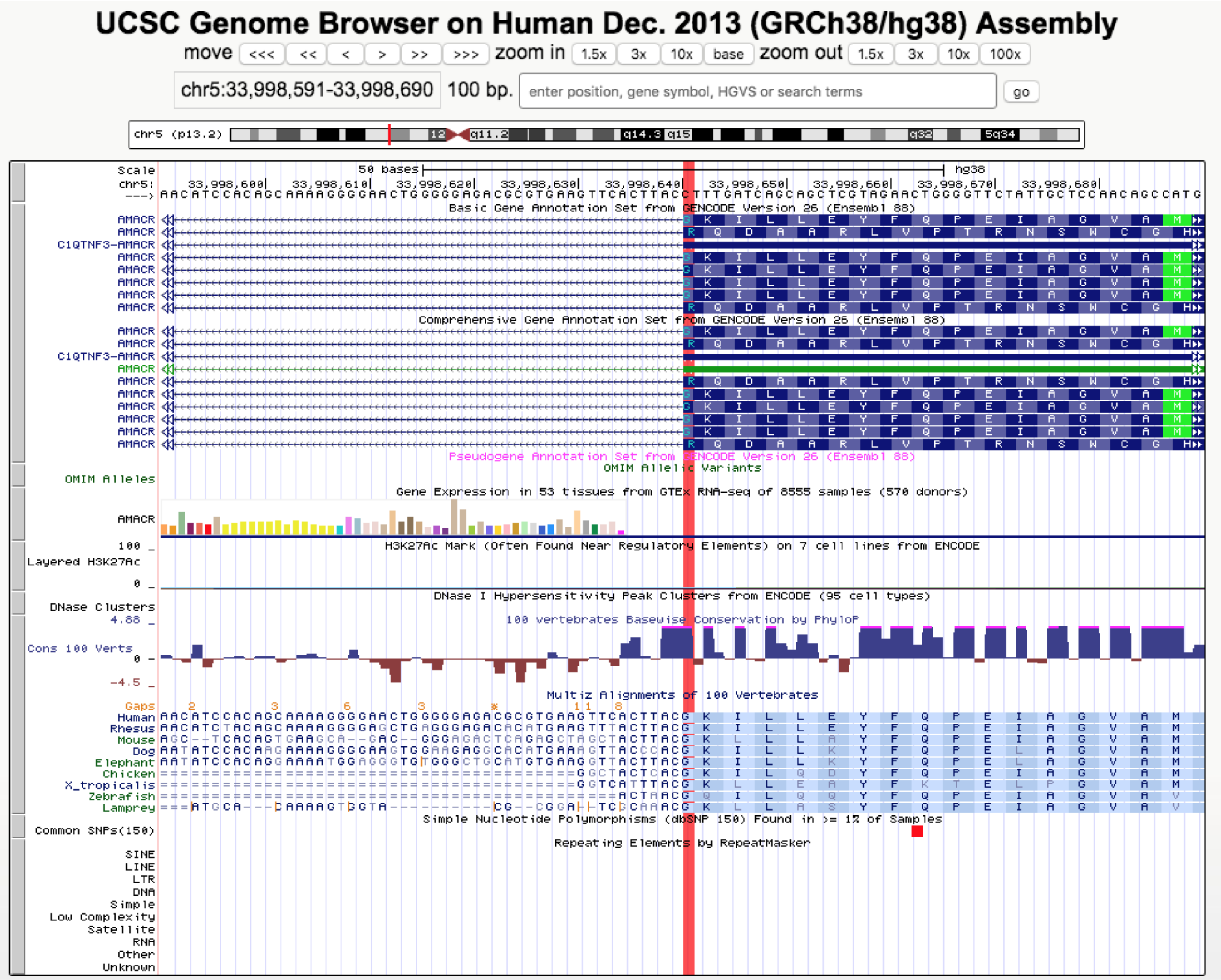


**Supplementary Figure 7.** A UCSC genome browser showing the 5' breakpoint (highlighted) for the two missed epitopes from the ENSG00000273294-ENSG00000164182 fusion. The 5' partner was reported as the readthrough transcript C1QTNF3-AMACR but the breakpoint is in the 5' UTR for C1QTNF3-AMACR. The overlapping AMACR contains the epitope predicted by INTEGRATE-Neo.

#### 2.2 Supplementary Tables

**Supplementary Table 1.** The overall metrics for the 326 samples PRAD ProTECT run.This table includes all metrics captured for each file including input file sizes, and numbers of mutations called and accepted, peptides generated, and IARs predicted.

**Supplementary Table 2.** The ProTECT ranks for all 7 vaccine candidates tested by Carreno et.al. (1).

**Supplementary Table 3.** Ranked IARs predicted from all 8 samples described in Carreno et.al.(1).

**Supplementary Table 4.** Detailed reasons for rejecting INTEGRATE-Neo(2) predicted neoepitopes. Table A describes 4 breakpoints that were missed due to issues with the 5' partner, Table B describes a breakpoint that should have failed similar to the case in Table 1, but were rescued by the prediction of a second, overlapping breakpoint, and Table C describes the INTEGRATE-Neo- and ProTECT-predicted binding affinity for the epitopes rejected as poor binders.

#### 2.3 Supplementary Files

**Supplementary File 1:** A tarball containing all the results from running ProTECT on 326 samples in the TCGA PRAD cohort.

**Supplementary File 2:** A tarball containing all the results from running ProTECT on 8 Melanoma samples described by Carreno et.al. (1).

**Supplementary File 3:** A tarball containing all the results from running ProTECT on 83 samples from the TCGA PRAD cohort, described by Zhang et.al (2). Specifically, this contains the results from running ProTECT using the fusions and HLA calls described in Zhang et.al.
