## Supplemental Data 1 for "ProTECT – Prediction of T-cell Epitopes for Cancer Therapy": car_t_target_report.txt

Available clinical trials for prostate cancer with GTEX and TCGA median values
	Gene      GTEX      TCGA N    Observed  DOI for gene papers                     Clinical Trials

	PSMA      79.99     124.25    258.69    10.1158/0008-5472.CAN-05-0436           NCT03089203 
	PSCA      188.07    346.59    24.20     10.1186/1471-2407-14-30                 NCT02744287 

Based on the genes overexpressed in this cancer type, here's a list of clinical trials for other types of cancer
	Gene      GTEX      TCGA N    Observed  DOI for gene papers                     Clinical Trials  Cancer              
